# Fucosylated and non-fucosylated α2,3 sialosides were detected on the bovine mammary gland tissues

**DOI:** 10.1101/2024.07.29.605565

**Authors:** Takahiro Hiono, Tatsuru Morita, Keigo Kosenda, Yuki Otani, Osamu Ichii, Norikazu Isoda, Yoshihiro Sakoda

## Abstract

Receptors for high pathogenicity avian influenza viruses (HPAIVs) in the mammary glands of dairy cattle were detected using various recombinant hemagglutinins (rHAs). Results demonstrated the presence of fucosylated and non-fucosylated α2,3 sialosides, which were typically targeted by the HA of clade 2.3.4.4b HPAIVs.

---

H5 subtype high pathogenicity avian influenza viruses (HPAIVs) of the A/goose/Guangdong/1/1996 lineage, especially those belonging to clade 2.3.4.4b, have been widely spread in the Eurasian, African, and American continents (1). The global spread of viruses now substantially affects not only avian species, but also mammals. Since the most affected mammals are carnivores and scavengers, it has been speculated that these mammals are infected by the ingestion of birds infected with HPAIVs (2,3). To date, sustained transmission of HPAIVs among mammals has not been observed frequently, although this possibility has not been excluded or has even been suspected in some cases (4–6).

The latest report of an outbreak of HPAIV infections in dairy cattle in the United States unexpectedly sheds light on the novel tissue tropism of the viruses (7). Ruminants, including dairy cattle, are considered less susceptible to influenza A virus infections (8). Unlike other mammals, the virus shows a preference for the mammary glands rather than the respiratory tract (7). Interestingly, the virus spread rapidly among the herds, and direct transmission from the mammary glands to the mammary glands was speculated.

Since the report of HPAIV cases in dairy cattle in the United States, several studies have attempted to detect sialylated glycans in the dairy cattle mammary glands as receptor molecules for HPAIVs (9,10). These studies used plant-derived lectins to detect influenza virus receptors. The use of plant-derived lectins needs careful consideration because they also recognize glycans that cannot be receptors for HPAIVs (11). Nwosu et al. previously demonstrated different magnitudes of sialylation and fucosylation between human and cow milk oligosaccharides (12), suggesting the necessity of detecting various sialosides in bovine mammary glands. However, this was not addressed in previous studies. Considering the fact that contemporary clade 2.3.4.4 viruses showed altered receptor-binding specificity to recognize various α2,3 sialosides (2,13), here, we analyzed the detailed receptor distributions of bovine mammary gland, especially focusing on the modification of α2,3 sialosides.

## The Study

Mammary glands were sampled from three dairy Holsteins (one clinically healthy and two clinically diagnosed with *Trueperella pyogenes* infection by milk culture). These included samples from a normal mammary quarter of a healthy individual (n = 2), a normal mammary quarter of *T. pyogenes*-infected individual (n =1), or a quarter with *T. pyogenes* detection (n = 3). These tissues were obtained from a local slaughterhouse and processed into formalin-fixed paraffin-embedded sections. Hematoxylin and eosin staining was performed according to standard protocols. Recombinant hemagglutinins (rHAs) were expressed in HEK293S GnT (-/-) cells and used as molecular probes to detect specific glycan motifs containing sialic acid. These rHAs include those derived from A/Ezo red fox/Hokkaido/1/2022 (H5N1) (Fox/Hok/22; the virus belonging to the clade 2.3.4.4b and recognizing non-modified, fucosylated and sulfated α2,3 sialosides (2)), A/duck/Mongolia/54/2001(H5N2) (Dk/Mng/01; recognizing only non-modified α2,3 sialosides (14)), A/chicken/Ibaraki/1/2005 (H5N2) (Ck/Ibr/05; exclusively requiring the fucosylation and being not sensitive for the 6-O-sulfation of the α2,3 sialosides (14)), and A/chicken/Tainan/V156/1999 (H6N1) (Ck/Tn/99; exclusively requiring the 6-O-sulfation and being not sensitive for the fucosylation of the α2,3 sialosides (15)). These rHAs were precomplexed with anti-Strep-tag II monoclonal antibody (StrepMAB-Classic; IBA Lifesciences; 1.0 μg/mL) and biotinylated antibody (goat anti-mouse IgG; human ads-BIOT, 1:200 dilution) for 30 min at 4°C. After blocking with 10% normal goat serum, the sections were incubated overnight at 4°C with the complex of rHAs and antibodies. Sections were washed with phosphate buffer saline and incubated with streptavidin-horseradish peroxidase (Nichirei Biosciences) for 30 min at room temperature. Sections were incubated with DAB-H_2_O_2_ solution, counterstained with Mayer’s hematoxylin, dehydrated using a series of alcohols, and cleared with xylene. Tissue sections were pretreated with sialidase for 12 h to prove the specificity of the positive signals.

As we collected specimens from retired milking cows, the tissue sections contained mammary lobules with expanded secretory mammary alveoli (lactation period), which were often filled with eosinophilic materials (most possibly milk; Figure 1A) and those with involuted alveoli (non-lactating period; Figure 1B). Lactiferous alveolar ducts were easily observed in the lobules during the non-lactating period. The rHAs derived from the contemporary clade 2.3.4.4b HPAIV, Fox/Hok/22, bound to the alveolar epithelia during lactation and non-lactation periods, as well as the epithelia lining alveolar ducts in the lobules and interlobular connective tissues (Figure 1C and D). This observation is consistent with the HPAIVs infection in dairy cattle mammary glands (7). These signals disappeared after sialidase treatment (Appendix 1, Figures 1A and B), confirming the specificity of the signals. Considering that only a limited number of amino acid differences were recognized between the HA of Fox/Hok/22 and dairy cattle isolates (Figure 2), these viruses share similar receptor-binding specificities.

**Figure 1.**
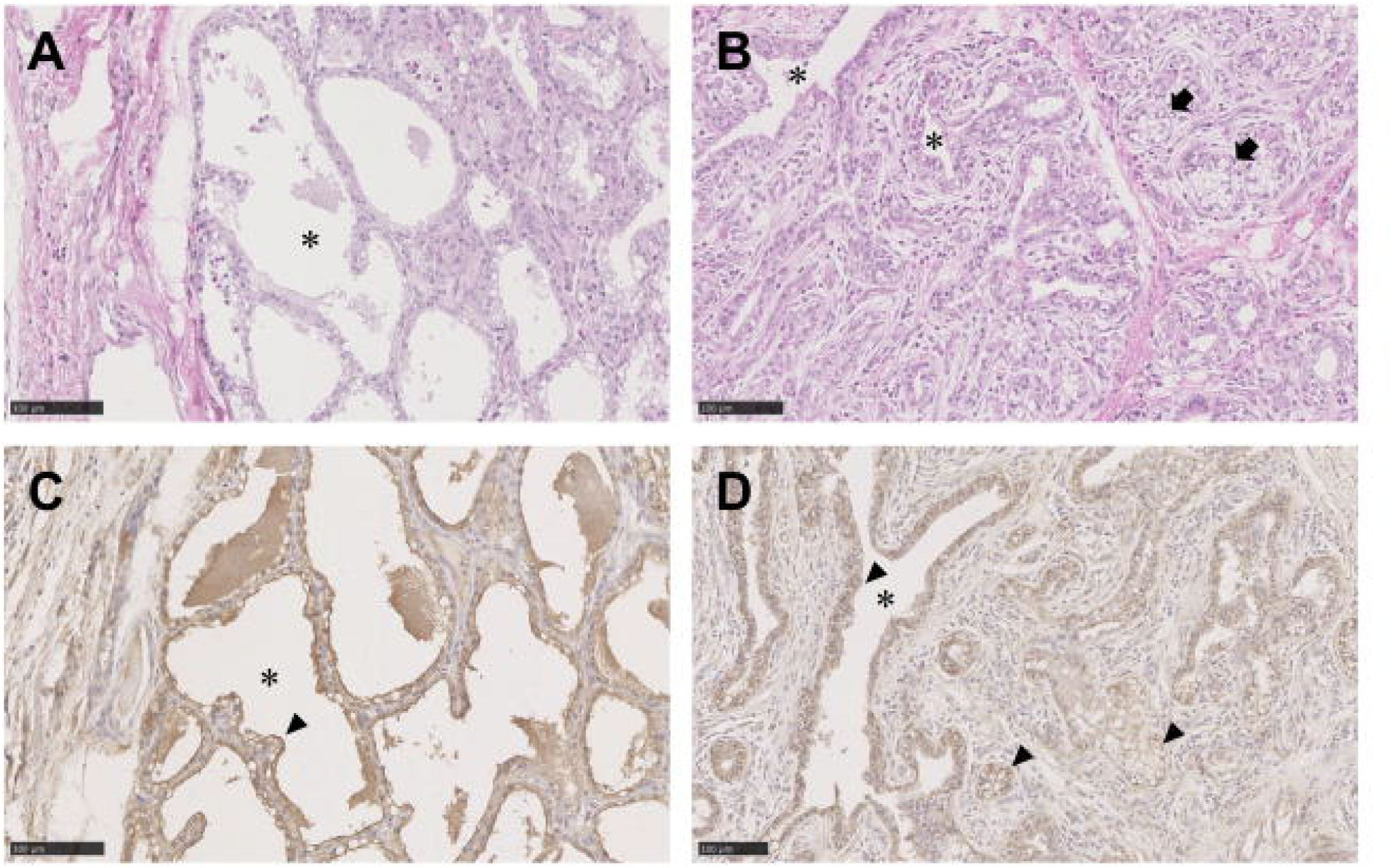
Receptors of highly pathogenic avian influenza viruses (HPAIVs) receptors in bovine mammary gland tissues. Hematoxylin and eosin staining of mammary lobules in lactation period (A) non-lactation period (B). Histochemical staining of mammary lobules in lactation period (C) non-lactation period (D) with recombinant hemagglutinins (rHAs) from Fox/Hok/22. Asterisks indicate the rumen of secretory mammary alveoli and lactiferous alveolar ducts. Arrows indicate involuted alveoli. Arrowheads indicate typical positive signals. Scale bars: 100 μm.

**Figure 2.**
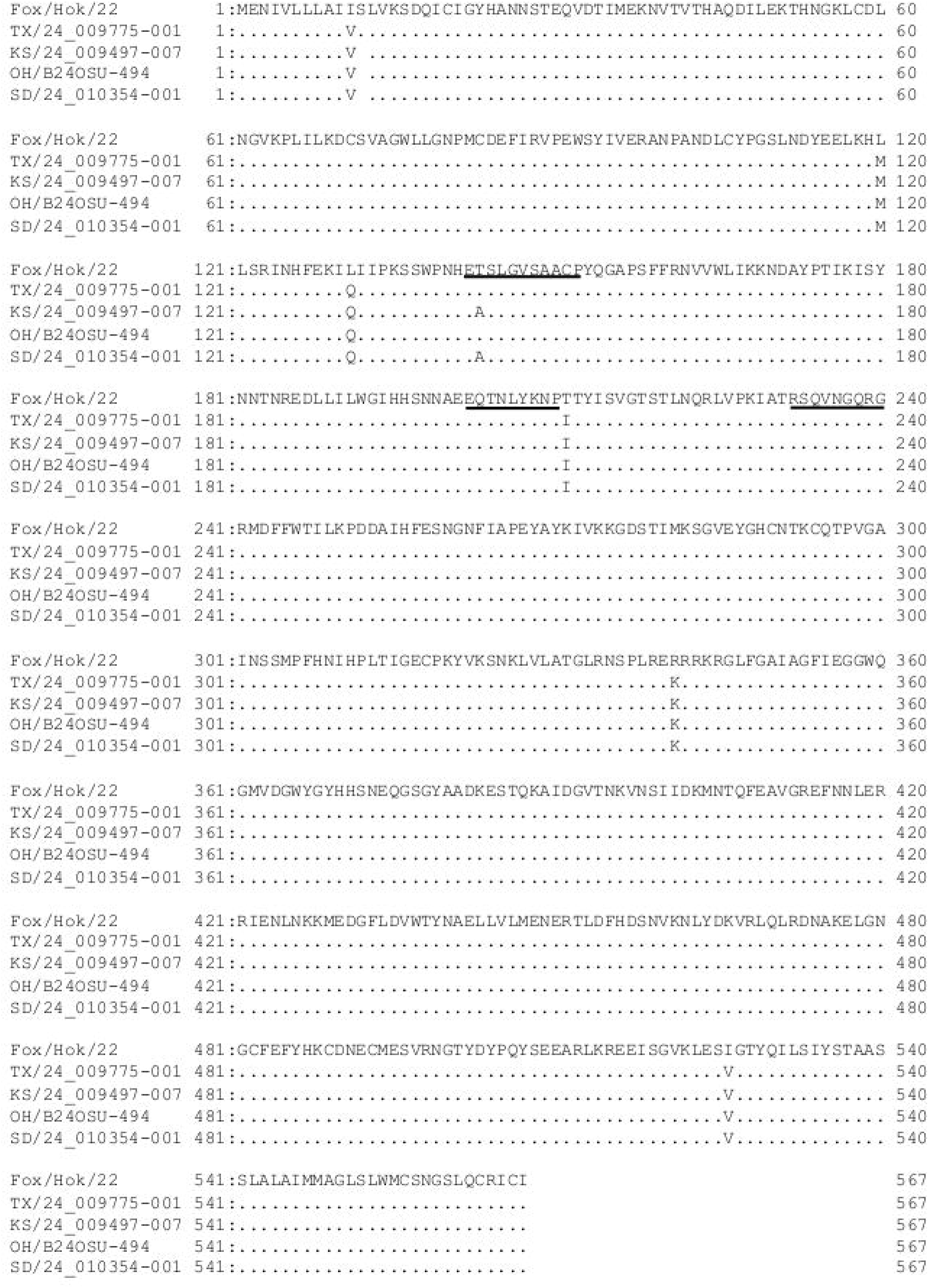
Amino acid sequence comparison among the hemagglutinins (HA) of Fox/Hok/22 and those of bovine H5N1 HPAIVs. Amino acid sequences of the HA were obtained from GISAID (https://gisaid.org/). Obtained sequences were aligned with Muscle and visualized using GENETX ver 16 (NIHON SERVER CORPORATION). Amino acids in 130-loop, 190-helix, and 220-loop were underlined. TX/24_009775-001: A/dairy cow/Texas/24_009775-001/2024 (H5N1). KS/24_009497-007: A/dairy cow/Kansas/24_009497-007/2024 (H5N1). OH/B24OSU-494: A/dairy cow/Ohio/B24OSU-494/2024 (H5N1). SD/24_010354-001: A/dairy cow/South Dakota/24_010354-001/2024 (H5N1).

Since the rHAs of Fox/Hok/22 recognizes a wide range of α2,3 sialosides, we further tried to analyze the detailed structure of these α2,3 sialosides using previously characterized rHAs (14,15). The binding of the rHAs from Dk/Mng/01 and Ck/Ibr/05 to the epithelia in mammary glands suggested the presence of both fucosylated and non-fucosylated α2,3 sialosides (Figure 3A-D). However, none of the epithelia were stained with the rHAs of Ck/Tn/99 (Figure 3E and F), suggesting that there was no sulfation of the antepenultimate GlcNAc for these receptors. These staining patterns suggested the presence of sialyl LacNAc and sialyl Lewis X glycoepitopes as α2,3 sialosides without 6-O-sulfation of antepenultimate GlcNAc in bovine mammary glands. These glycans are typically targeted by HA from clade 2.3.4.4b HPAIVs (2,13). Also, the results were well consistent with the report from Rios Carrasco et al. (11); the bovine mammary gland was stained with the rHAs derived from HPAIVs isolated in the mid-2000s (typically targeting only non-fucosylated α2,3 sialosides) and mid-2010s (typically targeting both fucosylated and non-fucosylated α2,3 sialosides (13)).

**Figure 3.**
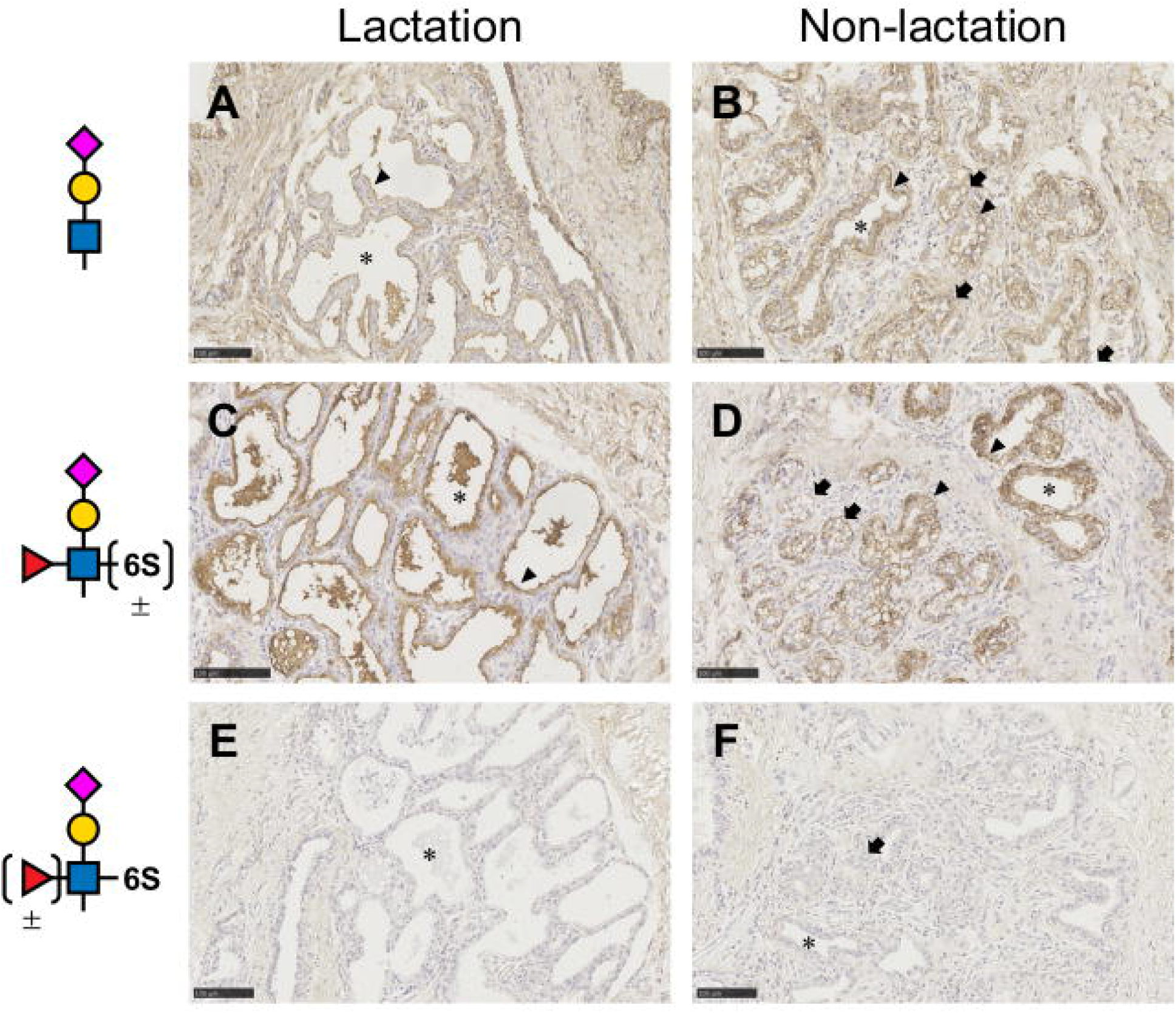
Distribution of α2,3 sialosides in bovine mammary gland tissues. Histochemical staining of mammary lobules in lactation period (A, C, E) non-lactation period (B, D, F) with rHA from Dk/Mng/01 (A, B), Ck/Ibr/05 (C, D), and Ck/Tn/99 (E, F). Symbols indicate the typical glycoepitopes detected by each of rHAs. Magenta diamonds: sialic acid. Yellow circles: Galactose. Blue rectangles: GlcNAc. Red triangle: fucose. 6S: 6-O-sulfation. Asterisks indicate the rumen of secretory mammary alveoli and lactiferous alveolar ducts. Arrows indicate involuted alveoli. Arrowheads indicate typical positive signals. Scale bars: 100 μm.

Interestingly, the subcellular localization of glycans was slightly different in the alveolar epithelia during the lactation and non-lactation periods. More specifically, the apical surface of the secretory mammary alveoli during lactation was strongly stained with rHAs from Ck/Ibr/05, whereas positive signals were observed in the basal membrane of the involuted alveoli (Figure 3C and D, arrowheads). A similar tendency was observed when rHAs from Fox/Hok/22 or Dk/Mng/01 were used, although this was not obvious when compared to rHAs from Ck/Ibr/05. No significant difference in the staining pattern was observed among individuals, even with and without *T. pyogenes* detection (Appendix 1, Figure 2).

## Conclusions

The present study clearly demonstrated the presence of fucosylated and non-fucosylated α2,3 sialosides in bovine mammary gland tissues. The receptor distribution was consistent with the well-known receptor-binding specificity of contemporary clade 2.3.4.4b HPAIVs. In addition, slight changes in the subcellular localization of glycans between the lactation and non-lactation periods may affect the susceptibility of dairy cattle to HPAIVs. Finally, experimental infection studies are essential to determine the susceptibility of dairy cattle to HPAIVs in different lactation stages.

## Acknowledgments

We would like to thank Nichiro Chikusan Co., Ltd., Mr. Yusuke Yamashita, and Ms. Natsumi Yamashita for supporting us with sample collection. We would also like to thank the authors and laboratories that identified and submitted the sequences to the GISAID EpiFlu database used in this study. All data submitters were contacted directly through the GISAID website (www.gisaid.org). This study was supported by the SHIONOGI Infectious Disease Research Promotion Foundation. This study was supported in part by the Japan Society for the Promotion of Science (JSPS) KAKENHI 24K09260 and the Japan Agency for Medical Research and Development (AMED; grant no. JP223fa627005).

## Authors Bio

Dr. Hiono is an assistant professor at Hokkaido University, Sapporo, Hokkaido, Japan. His research interests include elucidating the molecular basis of virus host range and pathogenesis using glycosicentific approaches. Mr. Morita is an undergraduate student at Hokkaido University, Sapporo, Hokkaido, Japan. His primary research focuses on the host range of influenza A virus.

## Figure legends

**Appendix Figure 1**. Histochemical staining of bovine mammary gland tissues after sialidase treatment. Histochemical staining of mammary lobules during the lactation (A, C, E, G) and non-lactation periods (B, D, F, H) with recombinant hemagglutinins (rHAs) from Fox/Hok/22 (A, B), Dk/Mng/01 (C, D), Ck/Ibr/05 (E, F), and Ck/Tn/99 (G, H) mice. Asterisks indicate the rumen of secretory mammary alveoli and lactiferous alveolar ducts. The arrows indicate involuted alveoli. Scale bars: 100 μm.

**Appendix Figure 2**. Histochemical staining of bovine mammary gland tissues with *Trueperella pyogenes*. Histochemical staining of mammary lobules during the lactation (A, C, E, G) and non-lactation periods (B, D, F, H) with recombinant hemagglutinins (rHAs) from Fox/Hok/22 (A, B), Dk/Mng/01 (C, D), Ck/Ibr/05 (E, F), and Ck/Tn/99 (G, H) mice. Asterisks indicate the rumen of secretory mammary alveoli and lactiferous alveolar ducts. The arrowheads indicate typical positive signals. The arrows indicate involuted alveoli. Scale bars: 100 μm.

