## Appendix Figure 2 for "Fucosylated and non-fucosylated α2,3 sialosides were detected on the bovine mammary gland tissues"

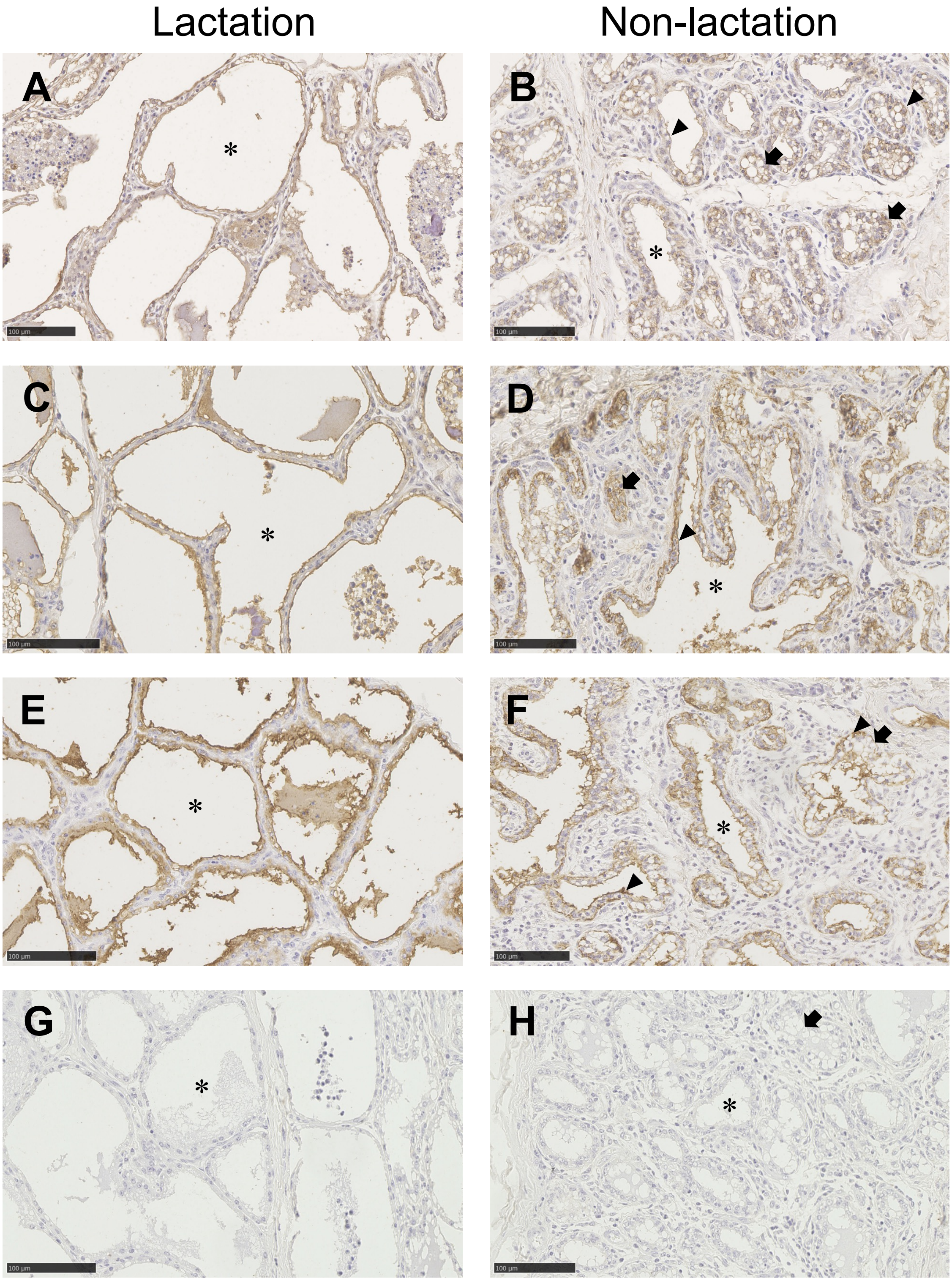


**Appendix Figure 2.** Histochemical staining of bovine mammary gland tissues with *Trueperella pyogenes*. Histochemical staining of mammary lobules during the lactation (A, C, E, G) and non-lactation periods (B, D, F, H) with recombinant hemagglutinins (rHAs) from Fox/Hok/22 (A, B), Dk/Mng/01 (C, D), Ck/Ibr/05 (E, F), and Ck/Tn/99 (G, H) mice. Asterisks indicate the rumen of secretory mammary alveoli and lactiferous alveolar ducts. The arrowheads indicate typical positive signals. The arrows indicate involuted alveoli. Scale bars: 100 µm.
